# Circular RNAs are Associated with Floral Fate Acquisition in Soybean Shoot Apical Meristem

**DOI:** 10.1101/2022.10.26.513951

**Authors:** Saeid Babaei, Mohan B. Singh, Prem L Bhalla

## Abstract

Soybean (*Glycine max*), a major oilseed and protein source, requires a short-day photoperiod for floral induction. Though key transcription factors controlling flowering have been identified, the role of the non-coding (dark) genome is limited. circular RNAs (circRNAs) recently emerged as a novel class of RNAs with critical regulatory functions. However, a study on circRNAs during the floral transition of a crop plant is lacking. We investigated the expression and potential function of circRNAs in floral fate acquisition by soybean shoot apical meristem in response to short-day treatment. Using deep sequencing and *in-silico* analysis, we denoted 384 circRNAs, with 129 exhibiting short-day treatment-specific expression patterns. We also identified 38 circRNAs with predicted binding sites for miRNAs that could affect the expression of diverse downstream genes through the circRNA-miRNA-mRNA network. Notably, four different circRNAs with potential binding sites for an important microRNA module regulating developmental phase transition in plants, miR156 and miR172, were identified. We also identified circRNAs arising from hormonal signaling pathway genes, especially abscisic acid, and auxin, suggesting an intricate network leading to floral transition. This study highlights the gene regulatory complexity during the vegetative to reproductive transition and paves the way to unlock floral transition in a crop plant.

**Highlight:** A new class of regulatory RNAs, circular RNAs, modulate floral transition in a crop plant, soybean, by regulating hormonal pathways and post-transcriptional processes.

## Introduction

Soybean is an economically important crop as it is the primary oilseed source and high-quality plant protein for the human diet and livestock feed. Global soybean production has increased about 13-fold from 1961 to 2017, mainly due to the increased cultivation of arable land (Liu et al., 2020). While the natural resources are limited, the demand for soybean and soybean products is rising. Thus, there is an urgent need to use modern crop breeding techniques to develop high-yielding soybean cultivars. Floral transition, or the switch from vegetative to reproductive growth, is a critical developmental stage in a plant life cycle, which ensures reproductive success and seed yield (Liew et al., 2014). Further, soybean with paleopolyploid genome arising from two whole-genome duplication events contains multiple copies of flowering genes. Understanding the molecular mechanisms is the first essential step for developing high-yielding crops and ensuring food security for the increasing world population, especially in changing climate.

The floral transition occurs in the shoot apical meristem (SAM), a multicellular dome-shaped structure located at the shoot apex. It contains a small population of undifferentiated and pluripotent stem cells. Endogenous signals and environmental factors regulate SAM growth and differentiation into organs such as stems, leaves, and flowers (Fouracre and Poethig, 2020). Photoperiod or day length is a critical environmental factor in regulating SAM development by extending the vegetative growth or inducing floral transition (Andrés and Coupland, 2012; Nakamichi, 2015). In Arabidopsis, a facultative long-day plant, environmental cues (temperature and photoperiod) integrate endogenous signals (phytohormones and plant age) to regulate floral pathway integrator genes such as FLOWERING LOCUS T (FT) and SUPPRESSOR OF OVEREXPRESSION OF CONSTANS 1 (SOC1). Regulation of these integrators then activates floral meristem identity genes such as APETALA1 (AP1), FRUITFULL (FUL) and CAULIFLOWER (CAL) which leads to SAM floral transition and flowering (Parcy, 2004). Photoperiod is also a critical factor that determines soybean flowering and seed yield. Soybean cultivars are classified into maturity groups based on the photoperiod requirement. Soybean is a typical short-day plant, i.e., long-day photoperiod extends the duration of vegetative growth, and short days promote floral transition. The transition of soybean SAM from vegetative to reproductive is visible after six short days of treatment (Wong et al., 2009b). At the molecular level, short-day treated soybean SAM displayed significant changes in gene expression profile, mostly in genes related to several members of the MADS-box transcription factor family and genes associated with hormones such as auxin, abscisic acid, and jasmonic acid (Haerizadeh et al., 2009; Wong et al., 2009a). Studies also reported the expression and potential function of non-coding RNAs such as microRNAs (miRNAs) and long non-coding RNAs (lncRNAs) in soybean SAM (Wong et al., 2011; Golicz et al., 2018). For example, the expression of 277 lncRNAs associated with floral transition observed in short-day treated soybean SAM (Golicz et al., 2018) suggests the role of regulatory RNAs in flowering. However, a study on circular RNAs is lacking.

CircRNAs are a distinct class of regulatory molecules, adding another layer of gene expression complexity in many eukaryotes (Liu and Chen, 2022). CircRNAs are a form of non-canonical RNA molecules that splicing machinery generates from precursor mRNA using a special kind of splicing termed back-splicing. Through back-splicing, a downstream 3’ splice site joins an upstream 5’ splice site to form a covalently closed circle RNA with no 5’ or 3’ polarities (Chen, 2020). Although most circRNAs have relatively low expression compared with their linear counterparts, they can play important roles in multiple biological processes in a tissue- or developmental stage-specific manner (Lasda and Parker, 2014; Wilusz, 2018; Babaei et al., 2021). Whilst the role of circRNAs is becoming increasingly apparent in human development, and diseases (Lee et al., 2019), very little is known about their role in plants. Recent studies in plants suggested that circRNAs are closely associated with growth, development and response to external stimuli (Zhao et al., 2019; Zhang et al., 2020).

To advance our knowledge of circRNAs during floral transition in soybean, we used micro-dissected SAM after short-day treatments in this study. Using four different bioinformatics prediction tools, we identified 384 circRNAs expressed in SAM at four-time points of short-day treatment. Further, the back-splicing junction of 26 selected circRNAs was experimentally validated. Differential gene expression analysis and functional enrichment annotation highlighted the function of circRNAs during the floral transition, especially circRNA production from genes related to hormonal signalling pathways, abscisic acid, and auxin. Further, miRNA–circRNA interactions highlighted the potential role of circRNAs as competing-endogenous RNAs (ceRNAs) in post-transcriptional gene regulation.

## Materials and Methods

### Plant Materials, and Sequencing

Soybean plants (*Glycine max*, cultivar Bragg) were grown under controlled conditions with a constant temperature of 25°C, 400 μmol m^−2^ s^−1^ light intensity, and 70% humidity. The plants were initially grown for 10 days in the long-day (LD) photoperiod (16 h light/8 h dark), and then transferred to the short-day (SD) treatment (8 h light/16 h dark) for two, four, and six days (Figure 1) (Zhang et al., 2021). For sample collection, shoot apical meristem (SAM) was dissected from about 100 plants within hours for each time-point: SD0 (LD10), SD2, SD4, and SD6, under a dissecting microscope as described before (Wong et al., 2008). Three biological replicates for each time points were used. Dissected SAM samples were immediately frozen in liquid nitrogen and stored at – 80 °C for later use.

**Figure 1.**
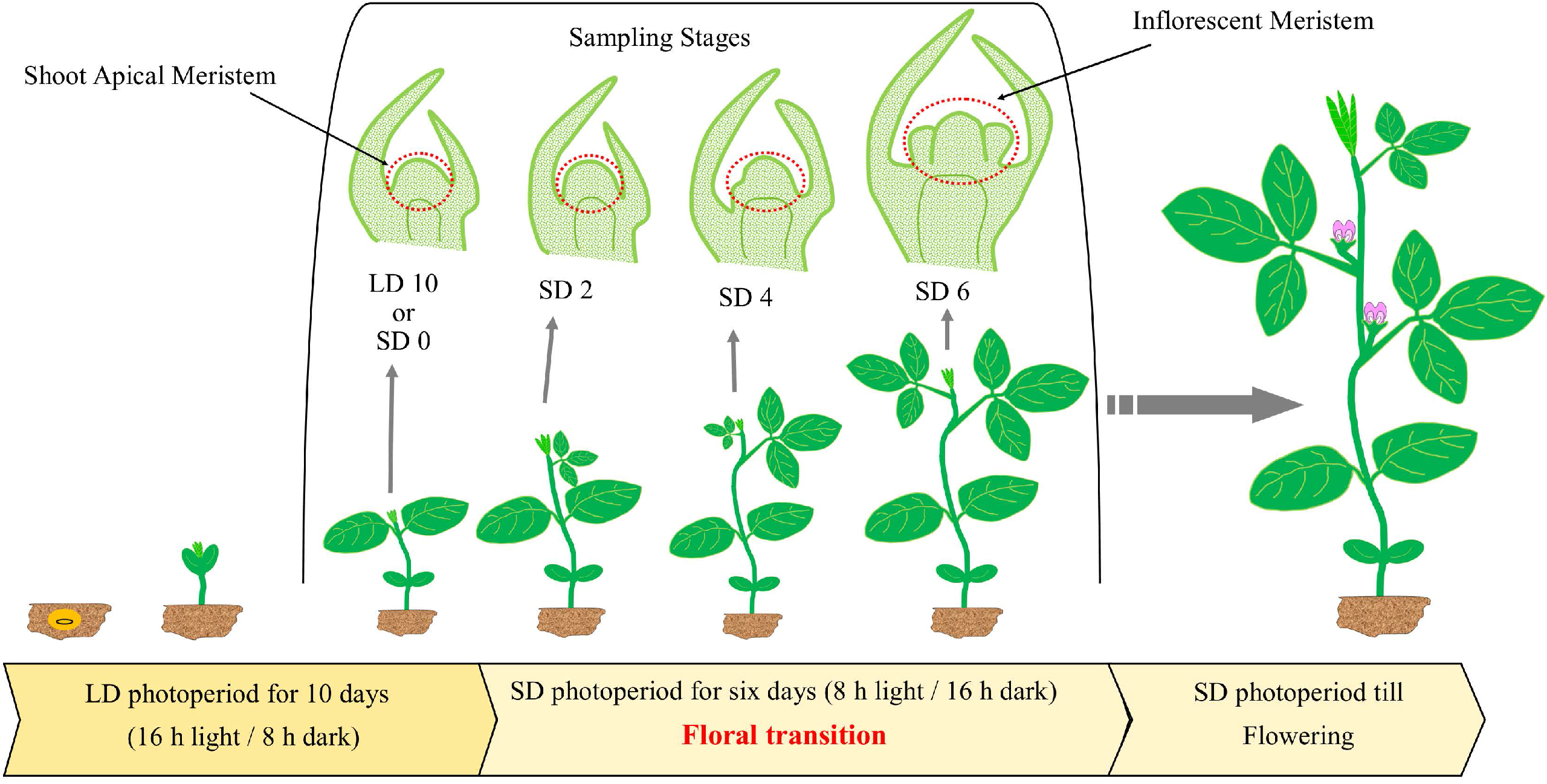
Schematic representation of sampling during soybean floral transition. Plants were grown for 10 days under long-day (LD) photoperiod (16 h light/8 h dark), and then transferred to short days (SD) (8 h light/16 h dark). In soybean, flowering occurs when plants expose to short day photoperiod, but floral transition generally induced within the first six days of short-day treatment. Red doted circles illustrate shoot apical meristem that were dissected from plants (SD 0, SD 2, SD 4, and SD 6). Stages of growth based on nomenclature in (Fehr and Caviness, 1977): VE: emergence, VC: unrolled unifoliolate leaves, V1: first trifoliolate, V2: second trifoliolate, and R1: beginning flowering.

Total RNA was isolated from SAM samples using the mirVana™ miRNA Isolation Kit (Thermo-Fisher; Part Numbers AM1560, AM1561, Carlsbad, CA, USA) according to the manufacturer’s instructions. TURBO™ DNase (Ambion, Carlsbad, CA, USA) was used to remove the remaining DNA from isolated RNA samples. The clean and high-quality RNA samples were used for circRNA sequencing (BGI Group, Hong Kong). The RNA-seq libraries were generated following RNase R treatment to remove linear RNAs. Paired-end read (100 bp) sequencing was performed using DNBseq Eukaryotic Transcriptome resequencing kit on DNBseq™ platform. The sequencing data were deposited to NCBI’s Sequence Read Archive (SRA) under accession number PRJNA893233.

### Identification, Characterization, and Differential Expression of Circular RNAs

To identify the expression of circRNAs in soybean SAM, we first mapped our RNA sequencing data against *Glycine max* reference genome (Wm82.a4.v1, Phytozome, https://phytozome-next.jgi.doe.gov/) (Goodstein et al., 2012; Valliyodan et al., 2019) using BWA (v0.7.17, mem-T 19) (Li and Durbin, 2009), Bowtie2 (v2.3.5.1) (Langmead and Salzberg, 2012), STAR (v2.7.5a) (Dobin et al., 2013), and CircMiner (v0.4.5) (Asghari et al., 2020) with their default parameters. Mapping results were then analyzed with four different circRNA detection tools: CIRI2 (v2.0.6) (Gao et al., 2018), find_circ (v1.2) (Memczak et al., 2013), CIRCexplorer2 (v2.3.8) (Zhang et al., 2016), and CircMiner (v0.4.5). After removing repeated circRNAs identified by two or more detection tools, a list of unique circRNA candidates with at least two supporting reads was generated for downstream analysis.

Using command lines from Bedtools (v2.30.0) (Quinlan and Hall, 2010), Bedops (v2.4.40) (Neph et al., 2012), and Genome-Tools (v1.6.2) (Gremme et al., 2013), we extracted genomic features and properties of our identified circRNAs based on the corresponding genome annotation file obtained from Soybase (https://soybase.org/genomeannotation/) (Grant et al., 2010). For comparison analysis, we used the aforementioned tools to create a list of probable linearly spliced exons as a control. The control list was created as follows: (i) a total of 10,000 genes (with the exons from only one mRNA isoform for each gene) were randomly selected from the soybean annotation file; all the circular producing genes identified in this study were removed from the selected genes; (ii) a diverse combination of exons from one exon up to six consecutive exons were extracted from the selected genes and saved as a list; (iii) similar to the portion of exon count in circRNAs found in this study (e.g., circRNAs that contain one exon, two exons, etc.), controls were randomly selected from the previous step list (Table S1).

Basic Local Alignment Search Tool (BLAST, v. 2.9.0) (Altschul et al., 1990) was used to find complementary elements in flanking introns of circRNAs and control exons, and RNAFold (Gruber et al., 2008) was used to compute the minimum free energy of the identified complementary sequences.

The R-Bioconductor package Noiseq (v2.28.0) (Tarazona et al., 2011) was used to quantify the expression level and to detect the differential expression pattern of circRNAs between samples. Trimmed Mean of M values (TMM) was used as the normalization method. A circRNA was considered differentially expressed if its q value was ≥ 0.8. Plots were generated using the R software (v4.0.4) (Team, 2013) with ggplot2 (v3.3.3) (Wickham, 2016), VennDiagram (v1.7.1) (Chen and Boutros, 2011), and gplots (v3.1.1) (Gregory R. Warnes, 2022) libraries.

### Analysis of circRNA–miRNA Interactions

To analyze the circRNA-miRNA interactions, we first downloaded the FASTA sequence of known miRNAs for soybean from miRbase database (Kozomara et al., 2019), and then we predicted the potential interactions between miRNAs and list of our identified differential expressed circRNAs using Targetfinder tool (v1.7) (https://github.com/carringtonlab/TargetFinder). We also predicted the mRNA targets of those miRNAs that had binding sites on circRNAs using Targetfinder. Cytoscape software (v3.8.2) (Shannon et al., 2003) was used to visualize the circRNA–miRNA interaction network.

### Functional Enrichment Analysis

Gene Ontology (GO) enrichment analysis of the host genes of differentially expressed circRNAs, and also the miRNA targets was carried out using topGO package (v2.42.0) (Alexa, 2021) with classic algorithm and fisher’s exact test. Then, we calculated the corrected *p*-value based on the Benjamini–Hochberg multiple testing correction (FDR < 0.05) to identify significant GO terms. To summarize the significant GO terms, we used ClueGO (v2.5.8) (Bindea et al., 2009) with its “preselected Functions”.

### Circular RNA Validation

We used RNase R treatment and PCR with divergent primers followed by Sanger sequencing to validate some of our identified circRNAs. Briefly, poly(A) was added to 10 µg of total RNA from each treatment using the Poly(A) Tailing Kit (Thermo Fisher Scientific AM1350) following the manufacturer’s protocol. Then, RNA samples were cleaned up using ethanol precipitation method (Green and Sambrook, 2020) and dissolved in nuclease-free water. The RNAs then were subjected to RNase R (Part number: E0111-20D1, Lucigen) treatment following the manufacturer’s instruction in a 20 µl reaction in the presence of lithium chloride containing buffer (Xiao and Wilusz, 2019). The RNA samples were cleaned up using ethanol precipitation protocol and reverse transcribed into complementary DNA (cDNA) using SuperScript™ III Reverse Transcriptase (Invitrogen, Carlsbad, CA, USA) in the presence of random hexamers. The sequence of 30 identified circRNAs that were predicted to be involved in SAM development were used for designing divergent primers (Panda and Gorospe, 2018) using PerlPrimer (Marshall, 2004). The PCR cycles were as follows: initial step at 95 °C for 5 min; followed by 40 cycles at 95 °C for 30 S, optimized annealing temperature (Table S2) for 40 s, and 72 °C for 30 s; and then a final cycle at 72 °C for 5 min. Electrophoresis of agarose gel (1%) was used to visualize the PCR products, and Wizard® SV Gel and PCR Clean-Up System (Promega, Madison, WI, USA) was used to recover the desired bands from the agarose gel. The recovered amplicons were cloned into the pJET1.2/blunt vector using the CloneJET PCR Cloning Kit (Thermo Scientific, Vilnius, Lithuania) and subjected to Sanger sequencing to validate the back-spliced junction sites.

## Results and Discussion

### Identification and characterization of 384 circRNAs in soybean SAM

To investigate the potential roles of circRNAs during floral fate acquisition by soybean SAM, circRNA-enriched RNA sequencing libraries (RNase R treated libraries for removing linear RNAs) were constructed for four short-day (SD) conditions: SD 0, SD 2, SD 4, and SD 6 with three biological replicates. RNA sequencing experiment produced over 1.4 billion clean reads with Q20 of about 98% for all 12 biological samples (Table S3). Analysis of the RNA sequencing data with four circRNA detection tools identified a total of 2580 circRNAs in all treatments with the criteria that a circRNA must be present in three biological replicates and supported by at least two back-spliced reads (Table S4). A total of 384 unique circRNAs were obtained by combining common circRNAs identified in different treatments and their biological replicates (Table S5). When comparing the number of circRNAs found by different detection tools, 320, 306, 293, and 260 circRNAs were found by CIRI2, CircMiner, CIRCExplorer2, and find_circ software respectively, and 181 circRNAs were common between four identification tools (Figure 2A).

**Figure 2.**
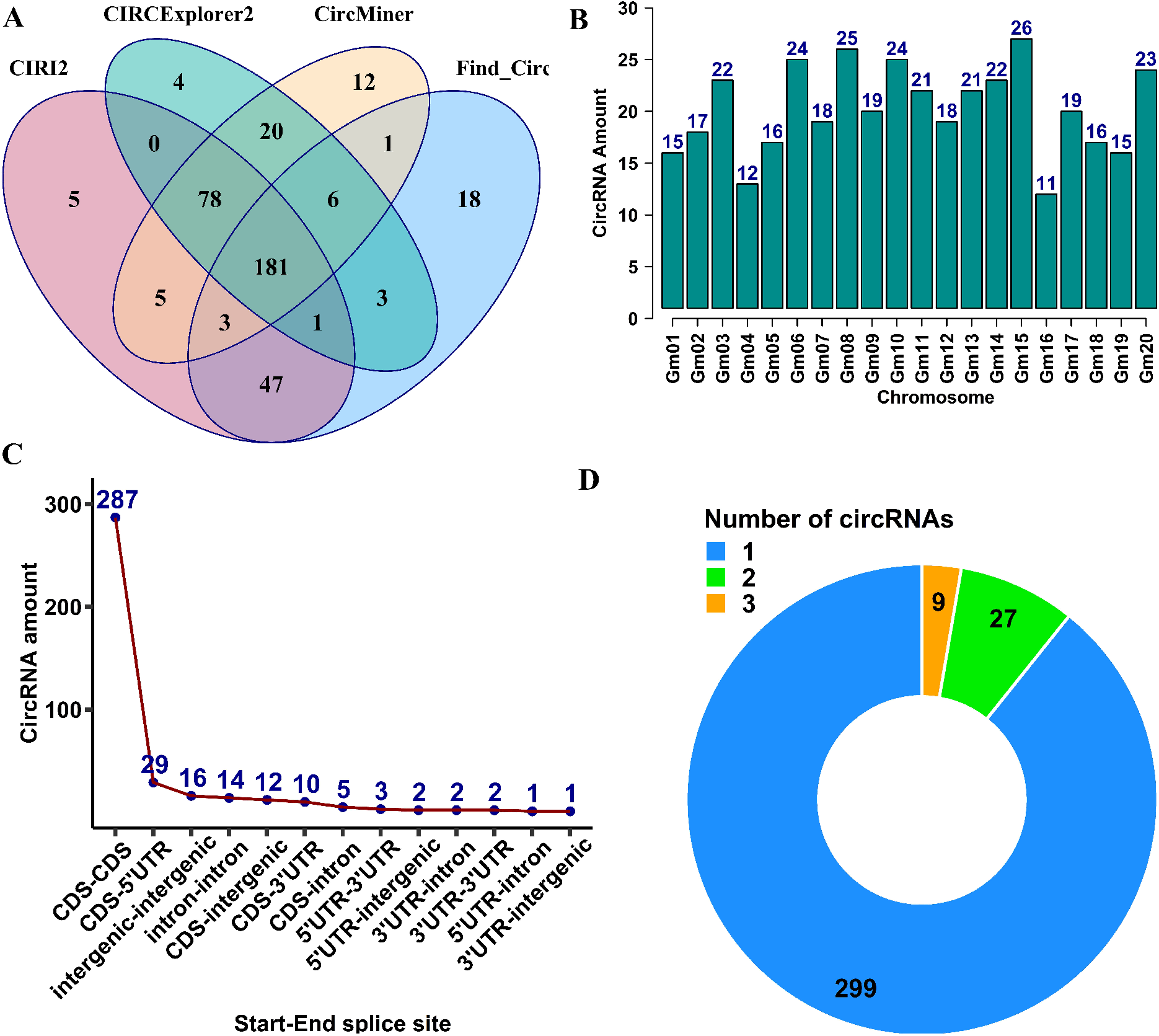
Features of identified circRNAs in soybean shoot apical meristem. (A) Venn diagram of unique and common circRNAs predicted by four detection tools. Of 384 identified circRNAs, 181 were common among identification tools. (B) Distribution of circRNAs on chromosomes. CircRNAs are distributed unevenly on different chromosomes. Chromosome 16 generated the most, and chromosome 11 generated the least amount of circRNAs. (C) The number of circRNAs generated from each genomic region. The vast majority of circRNAs overlapped entirely or partially with genic regions. (D) The parent genes of circRNAs produced different numbers of circular transcript isoforms. Of 335 circular producing genes, 299 generated only one circRNA isoform.

The distribution of circRNAs among chromosomes showed uneven transcription of circRNAs from different chromosomes; chromosome 15, with 26 circRNAs, generated the most, and chromosome 16, with 11 circRNAs, generated the least amount of circRNAs (Figure 2B). Moreover, the annotation analysis revealed that circRNAs could arise from diverse genomic regions, including Coding DNA Sequences (CDS), 5’ and 3’ Untranslated Regions (UTR), as well as intronic and intergenic regions (Figure 2C, Table S5). Among 384 identified circRNAs, 92.19% (354) of circRNAs contain at least one exon from coding sequences, 4.17% were intergenic, and 3.64% were from intronic regions (Figure 2C). While most of the circRNA-producing genes could generate only one isoform of circRNAs, 36 genes were able to produce two or three isoforms of circRNAs through alternative back-splicing (Figure 2D). We also noted ten circRNAs generated from two or three adjacent genes (Table S5).

### Experimental validation of circRNAs using Divergent primers and Sanger sequencing

To confirm the authenticity of the identified circRNAs, we selected 30 circRNAs whose parent genes or their miRNA targets were predicted to be involved in floral transition in soybean SAM. A set of divergent primers listed in Table S2 were designed for amplifying the back-splicing junction of each selected circRNA. Twenty-six circRNAs were successfully validated using PCR with divergent primers. We further checked the accuracy of the back-splicing sites by using 50% of amplified circRNA (13) to Sanger sequencing. The results verified the junction sequence of all the samples (Figure 3, and the Supplementary document). It is worth noting that all the validated circRNAs were made of only exons with introns spliced out.

**Figure 3.**
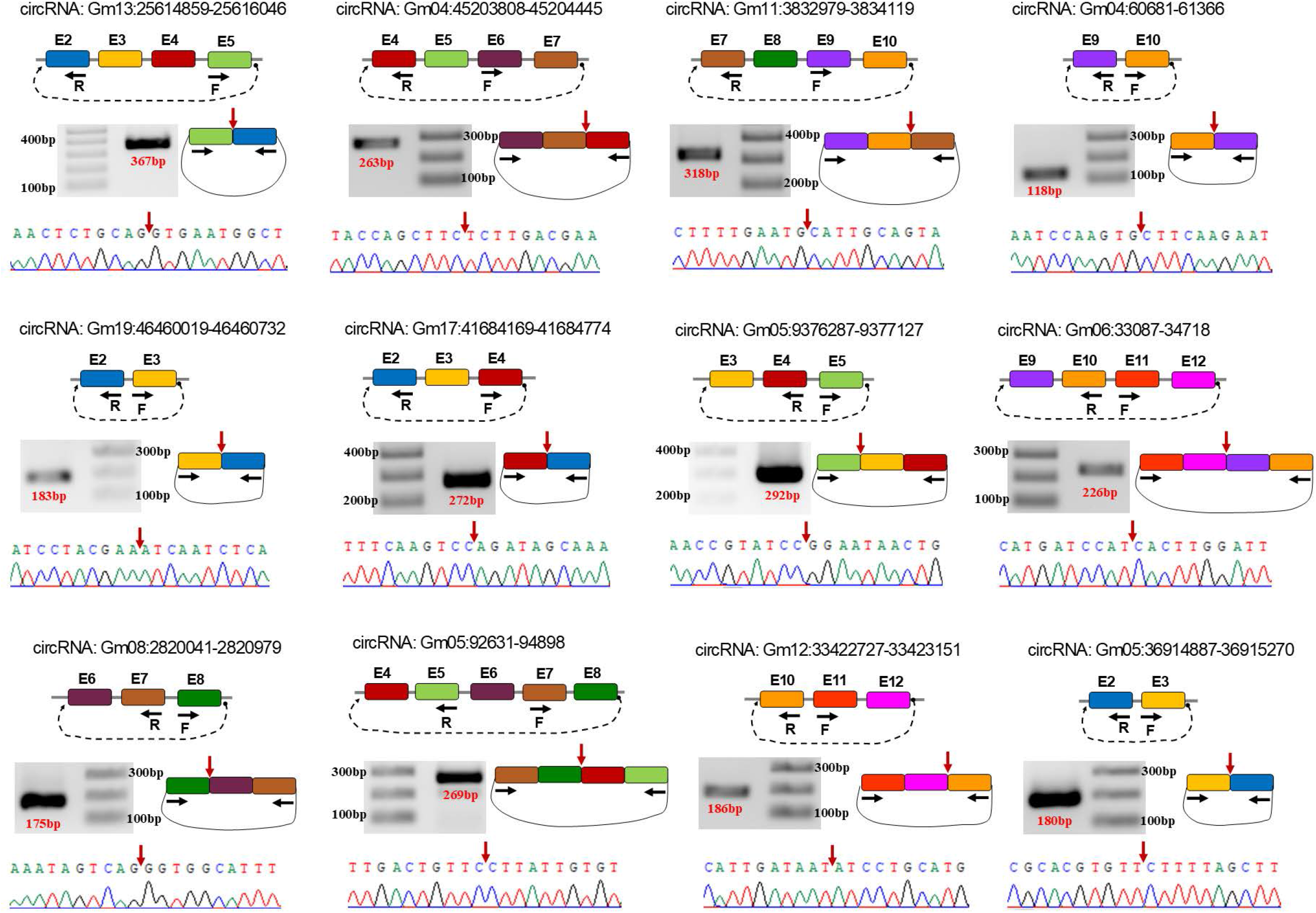
Validated circRNAs in soybean shoot apical meristem using divergent primers and Sanger sequencing. In each section, from top: circRNA ID; the exons (E) involved in circRNA; black arrows representing the position of forward (F) and reverse (R) primers; the amplified circRNA junction and its size on agarose gel (1% TAE buffer); red arrow representing the position of circRNA junction; confirmed circRNA junction sequence by Sanger sequencing.

### Exonic circRNAs contain multiple exons, and their overall length is under 1000 nucleotides

Exonic circRNAs in plants could contain several numbers of exons. However, they are usually composed of one to four exons in different plant species, such as soybean (78.2%) (Zhao et al., 2017b) and Arabidopsis (88.8%) (Philips et al., 2020). CircRNAs also tend to harbour their exons from the middle exons of their parent genes rather than the first or last exons in humans (Zhang et al., 2014) and plants. For example, in *Salvia miltiorrhiza*, the circRNAs generated from middle exons of their parent genes were 90.86%, 88.27%, and 85.46% in root, stem, and leaf tissues respectively (Jiang et al., 2021). The length of exons is also an important feature that allows exons to become circularized in eucaryotic cells (Chen, 2020). For instance, in humans, while an average exon length of about 350 bp was required for single-exon circularization, this number was about 120 bp for multiple-exon circRNAs (Zhang et al., 2014). This is true in some plant species, such as cotton (Zhao et al., 2017a) and cucumber (Zhu et al., 2019). Here our analysis revealed that exonic circRNAs were composed of one to 14 exons, but most (66.15%) contained two to four exons (Figure 4A). The exons in multiple-exon circRNAs mainly originated from the middle exons (84.89%) of their annotated genes rather than the first or last exons; nine circRNAs were also found to be originated from the single-exon genes (Figure 4B). The length distribution of the identified circRNAs was between 100 to 1500 nucleotides (nt). When considering the length of exons in exonic circRNAs, the length distribution was mainly between 100 to 900 nt, with only a few circRNAs longer than this range (Figure 4C). We also compared the length of exons between single-exon circRNAs and multiple-exon circRNAs. The results showed that single-exon circRNAs contained exons that were significantly longer than exons in multiple-exon circRNAs (Figure 4D).

**Figure 4.**
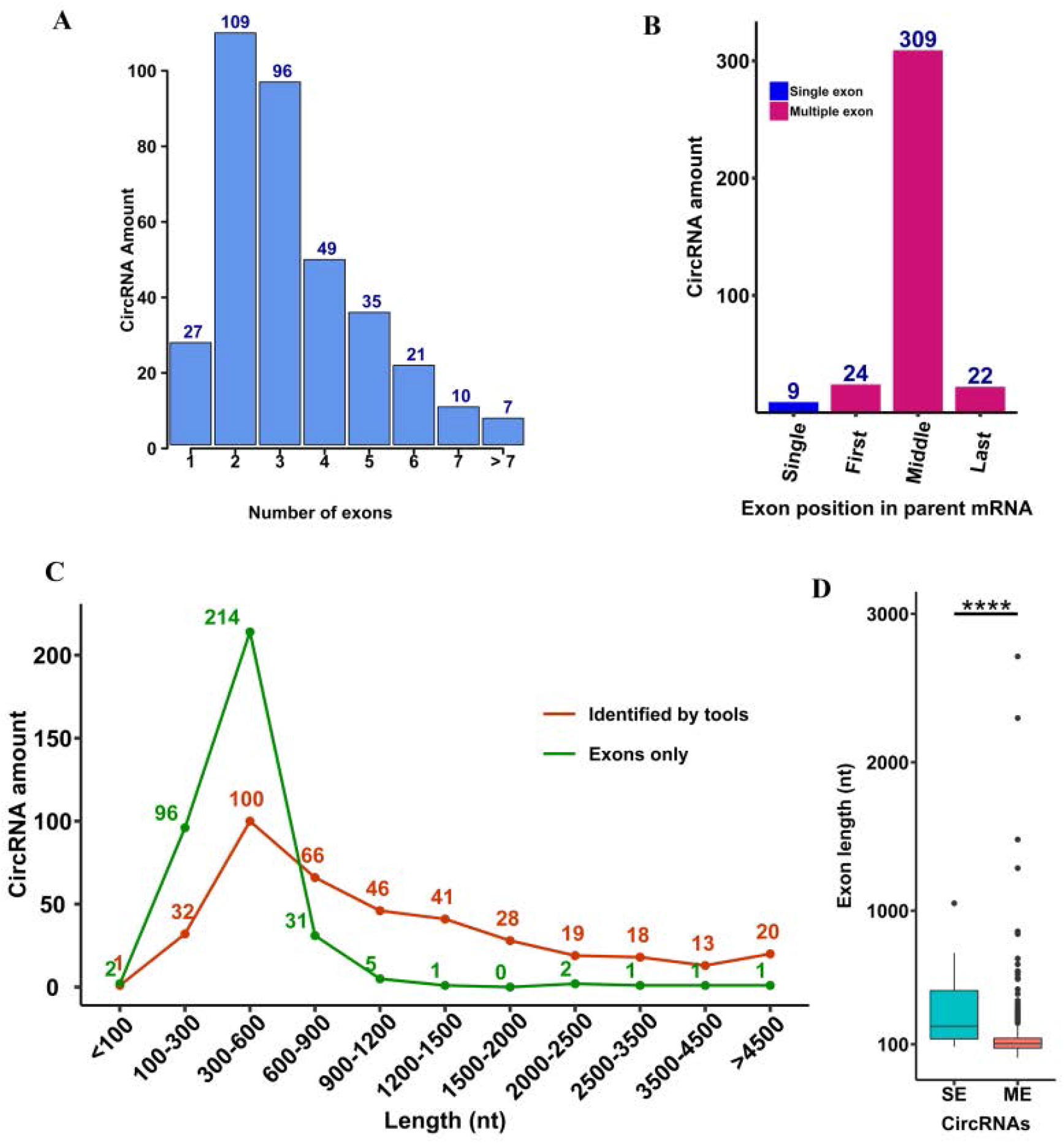
Features of exonic circRNAs expressed in soybean shoot apical meristem. (A) The number of exons in each circRNA. CircRNAs were mainly composed of two to four exons. (B) The position of exons in the parent gene of circRNAs. A large number of circRNAs originated from the middle exons of their parent genes. (C) Length distribution of circRNAs. Red colour: the length of circRNAs originally identified by detection tools. Green colour: the length of exonic circRNAs after removing introns. The peak length was between 300 to 600 nucleotides (nt). (D) Comparison of exon length between single-exon (SE) and multiple-exon (ME) circRNAs. SE circRNAs contain significantly longer exons compared with ME circRNAs (\*\*\*\**p* value = 2.1e^-9^, Wilcoxon rank-sum test).

### Reverse complementary sequences in the flanking introns could promote circRNA biogenesis in soybean SAM

Earlier studies have demonstrated that most circRNAs are flanked by longer than average introns with the existence of complementary sequences such as Alu elements in humans (Jeck et al., 2013; Zhang et al., 2014). Base pairing between these intronic elements, which could be as short as 30 to 40 nt, promotes the biogenesis of many circRNAs in mammals (Liang and Wilusz, 2014; Zhang et al., 2016). In plants, while circRNAs were found to be flanked by longer introns in species such as rice (Ye et al., 2015) and maize (Han et al., 2020), the amount of reverse complementary sequences was fewer compared to animals (Zhao et al., 2019). Nevertheless, very short complementary sequences ranging from 4 to 11 nucleotides near the splice sites of circRNAs might facilitate circRNA formation in Arabidopsis (Sun et al., 2016). Here we compared the length of introns flanking circRNAs with the length of introns flanking a list of simulated lists of control exons prepared as detailed under the materials and methods section. The results showed that circRNAs were flanked by significantly longer introns in their upstream and downstream regions (Figure 5A). The BLAST command line was used to scan upstream and downstream introns of circRNAs and control exons to check the presence of reverse complementary sequences (Table S6). The number of complementary bases and the best BLAST score (bitscore) were significantly higher in flanking introns of circRNAs compared with the control list (Figures 5B and 5C). As this higher statistical significance could be just the result of longer introns that flank circRNAs, statistical analysis was used to test the BLAST score against the length of flanking introns. The results revealed that in some ranges of intron length, especially from below 500 nt up to about 1000 nt, flanking introns of circRNAs contain a significantly higher amount of reverse complementary sequences compared with controls (Figure 5D). The comparison at longer intron size (above 5000 nt) was not possible as introns flanking controls were not in that size range. We also explored the presence of reverse complementary elements in the immediate 2 kb of the flanking sequences (not necessarily introns) of circRNAs and controls (Table S7). The result still displayed a higher BLAST score in flanking sequence of circRNAs, but with less degree than the flanking intron results (Figure 5E). To further investigate the flanking introns of circRNAs, the minimum free energy (MFE) of the identified reverse complementary sequences in the flanking introns of circRNAs and controls were compared. The results revealed significantly lower MFE in flanking introns of circRNAs which means stronger base-pairing between upstream and downstream reverse complementary elements (Figure 5F). All these results suggest that reverse complementary sequences in the flanking introns of circRNAs might play a role in circRNA biogenesis in soybean SAM.

**Figure 5.**
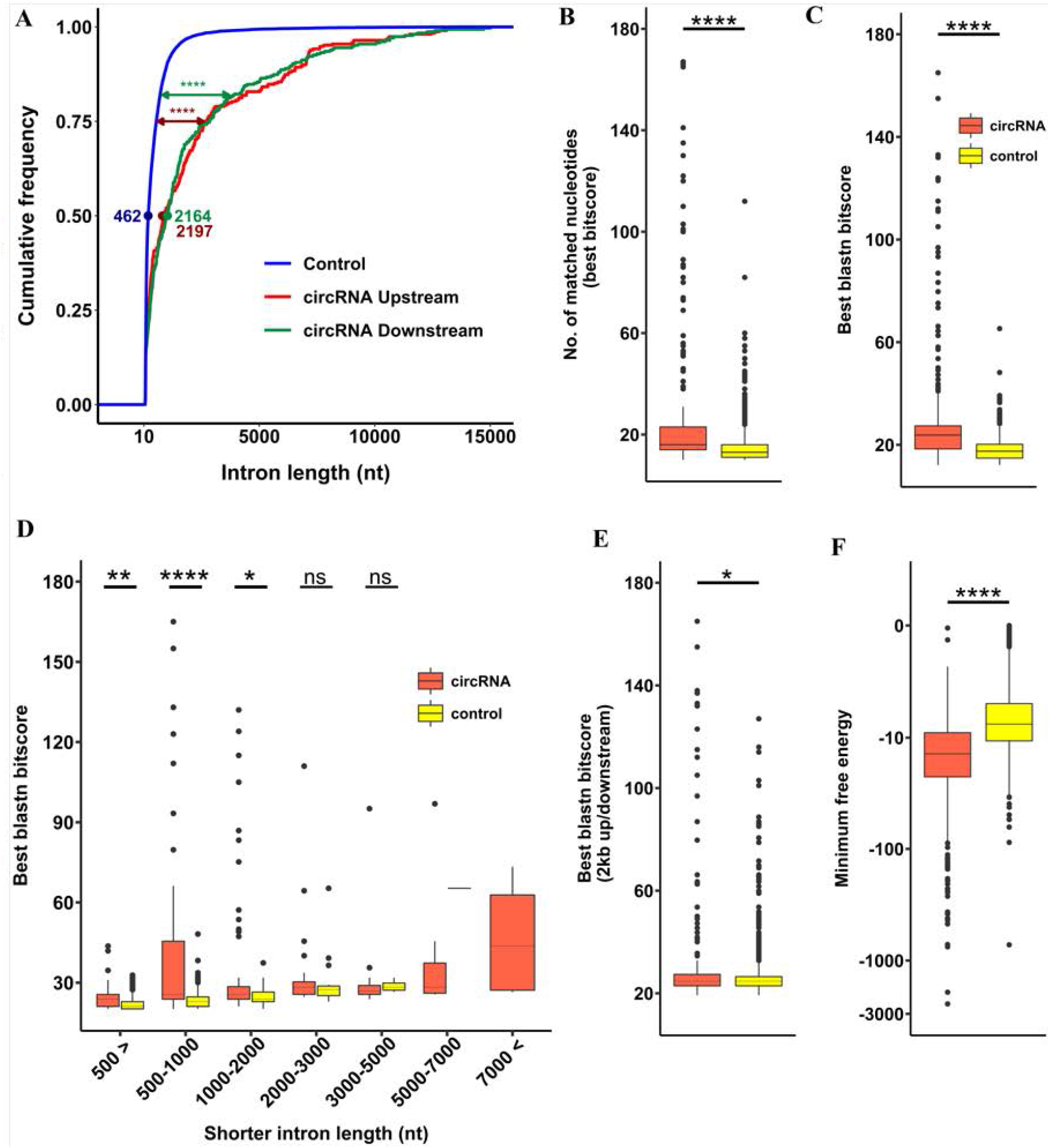
Features of introns flanking circRNAs. Wilcoxon rank-sum test was used as a statistical test in all sections. (A) Length distribution of upstream (red) and downstream (green) flanking intron of circRNAs compared with flanking intron of control exons (blue). Flanking introns of circRNAs were significantly longer than controls (\*\*\*\**p* value <2e^-16^). (B) The number of complementary nucleotides found between upstream and downstream introns of circRNAs and controls. Flanking introns of circRNAs contained much more complementary nucleotides than controls (****p value <2e-16). (C) Comparison of best BLAST scores between flanking introns of circRNAs and controls. Flanking introns of circRNAs had significantly higher scores which mean stronger alignment between identified complementary sequences (****p value <2e-16). (D) Length distribution of flanking introns and their BLAST score. Flanking introns of circRNAs had higher BLAST scores compared with controls, especially in the ranges below 500 up to 1000 nucleotides (**p-value 0.00613, ****p value 1.6e^-8^, *p-value 0.01314). (E) Comparison of best BLAST scores between flanking sequences of circRNAs and controls. Here the flanking sequences were selected from the immediate 2000 nucleotides upstream and downstream of circRNAs and controls. Flanking sequences of circRNAs had slightly higher scores than controls (*p-value 0.0048). (F) Minimum free energy of identified complementary sequences in the flanking intron of circRNAs and controls. Complementary sequences in the flanking intron of circRNAs had significantly lower minimum free energy, which means stronger base paring (****p value <2e-16). In sections B, C, D, and E, numbers above 180 on the vertical axis were used in the statistical test but not shown in the plots for graphical reasons.

### CircRNAs have distinct expression patterns in soybean SAM in response to short-day treatment

To explore the expression pattern of circRNAs during floral transition in soybean SAM, we compared the expression profile of circRNAs in each photoperiod time-point with its previous time-point (Table S8). The results revealed differential expression of 351 circRNAs, of which 129 were expressed explicitly in response to different photoperiod treatments (Figure 6A). The overall number of differentially expressed circRNAs tended to up-regulate following four short days and down-regulate after six short days (Figure 6B). However, when we visualized the expression pattern of 129 circRNAs specifically expressed in response to photoperiod treatment, we observed that in the early short-day treatment, more circRNAs showed down-regulated expression, and in short-day six, upregulation was more prevalent (Figure 6C). We also visualized the expression profile of circRNAs vs. different photoperiod treatments and their biological replicates using a clustering heatmap (Figure 6D). The expression analysis results indicated that circRNAs have a distinctive expression pattern in response to photoperiod treatment, suggesting their potential roles in floral transition of soybean SAM.

**Figure 6.**
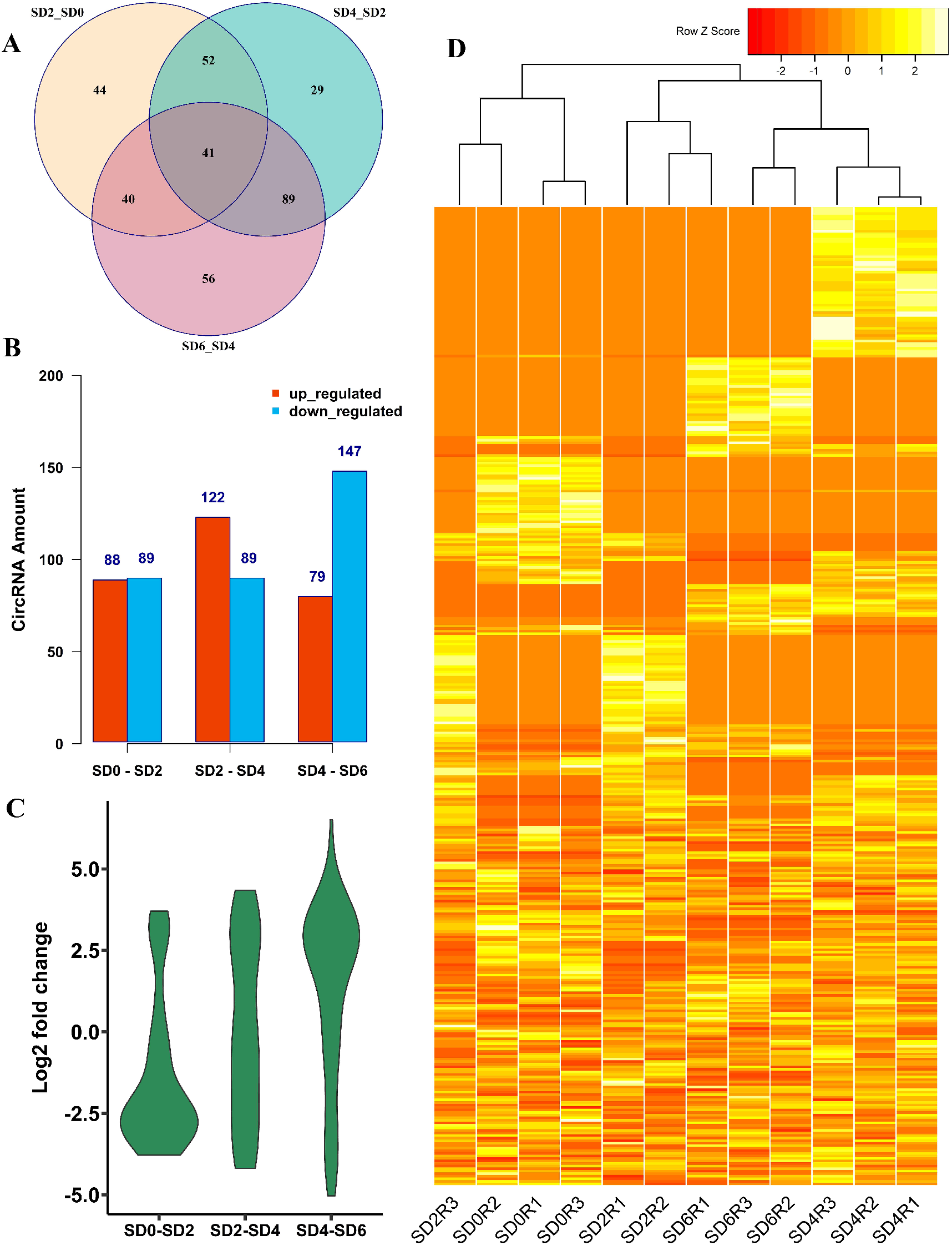
Expression pattern of circRNAs in soybean shoot apical meristem. (A) Venn diagram illustrates the number circRNAs with differential expression in response to short-day photoperiod treatment. Of 351 differentially expressed circRNAs, 129 showed treatment-specific expression patterns, while 41 were expressed commonly at all time-point of photoperiod treatment. (B) Histograms represent the number of up- or down-regulated circRNAs responding to photoperiod treatment. (C) The Violin plot illustrates the expression pattern of 129 treatment-specific circRNAs. The expression pattern of circRNAs changed when the plants were exposed to different time-point of photoperiod treatment. While the expression of treatment-specific circRNAs down-regulated when plants transferred to short-day photoperiod, the expression of circRNAs up-regulated after six short-day of photoperiod. (D) The heatmap shows the expression profile of 384 identified circRNAs in all samples with their biological replicates.

### Differentially expressed circRNAs contain one to four miRNA binding sites

To further evaluate the function of circRNAs in post-transcriptional gene regulation during floral transition in soybean SAM, we predicted the miRNA binding sites on differential expressed circRNAs (Table S9). The results revealed 38 circRNAs with putative binding sites for 39 miRNAs. Most circRNAs (29 out of 38) could target only one miRNA, and we found seven circRNAs that contained two potential miRNA binding sites. We also noted that circRNA Gm11:18737798-18738745 and Gm18:47348392-47380009 could target three and four miRNAs, respectively (Figure 7). Similarly, miRNAs could be targeted by a different number of circRNAs. Of 39 identified miRNAs, 33 had only one binding site on circRNAs, and six were targeted by two to four circRNAs (Figure 7). For example, Gma-miR156 and Gma-miR172 were the miRNAs that four circRNAs could target. These results suggest that circRNAs might be involved in gene regulation at the post-transcriptional level in soybean SAM.

**Figure 7.**
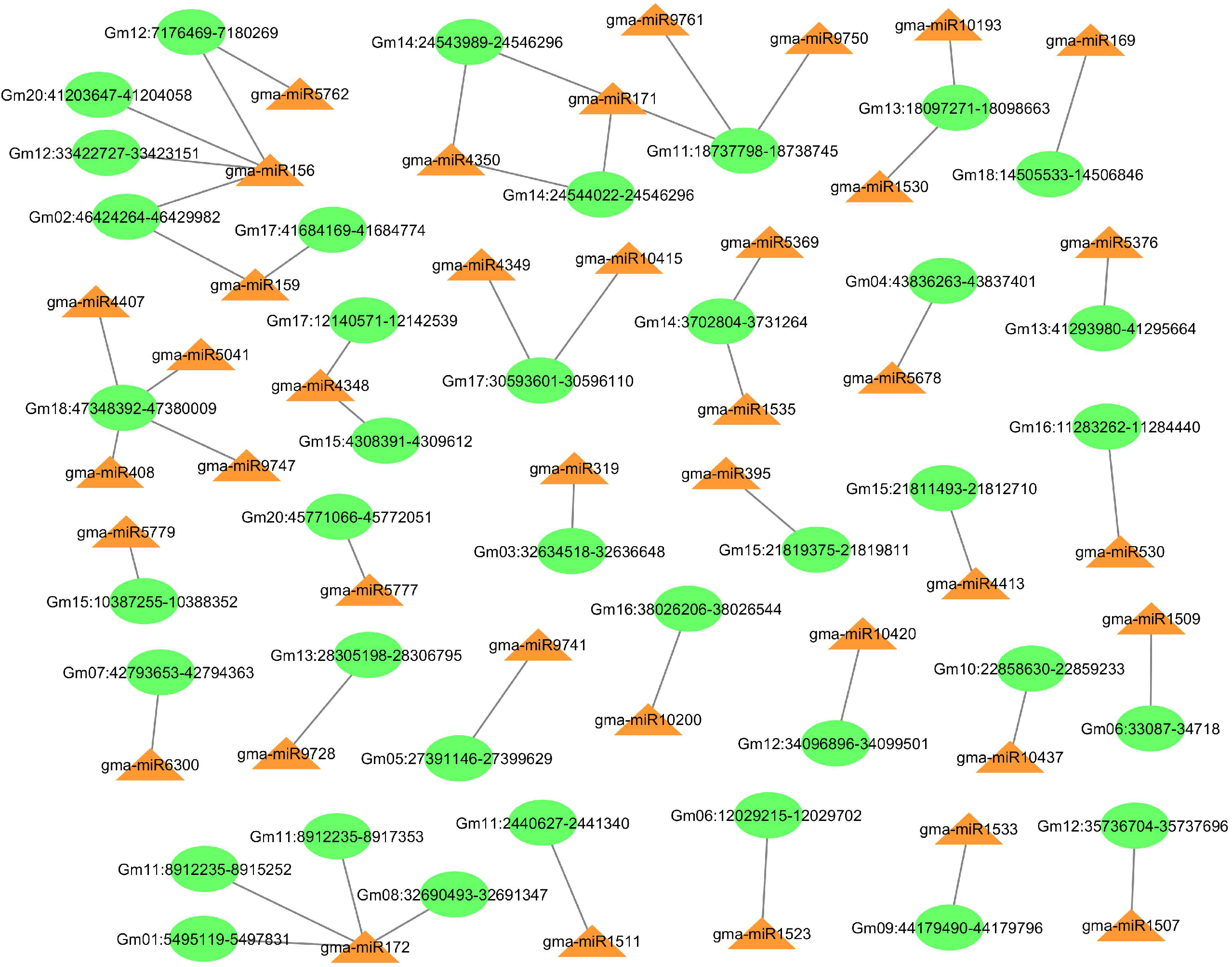
Predicted circRNA-miRNA interactions for differentially expressed circRNAs in soybean shoot apical meristem. CircRNAs could potentially contain one to four miRNA binding sites.

### CircRNAs function during floral transition through miRNA sponging

In our experiments, we further explored the target genes of well-known miRNAs, miR172 and miR156. MiR172 could target 164, and miR156 could target 199 downstream genes, mainly related to meristem development and flowering. For example, soybean genes Glyma.11G117300 and Glyma.11G117400, which are involved in the vegetative to the reproductive phase transition of the meristem, were able to produce an exonic circRNA from the genomic region Gm11:8912235-8915252 (circ-GmCCR2). Circ-GmCCR2 had downregulated expression in short-day treated plants and could potentially target miR172. Downregulation of circ-GmCCR2 can lead to upregulation of FLOWERING LOCUS T (FT) and initiation of floral transition as miR172 target genes such as Gm-AP2 and Gm-TOE1, which negatively regulate the expression of GmFT. We also noted two circRNAs with upregulated expression trends in short-day treated samples, which could further promote floral transition through sponging miRNA 156 (Figure 8). Previous studies in animals and humans have revealed that circRNAs can act as miRNA sponges suggesting their potential function as post-transcriptional gene expression regulators. Typical examples of such circRNAs are circSry, with 16 binding sites for miR-138 in mice testis and CDR1as has 70 binding sites for miR-7 in the mammalian brain (Hansen et al., 2013; Memczak et al., 2013). Studies have shown that when the expression of CDR1as is reduced in human cells, the level of mRNAs with binding sites for miR-7 decreases indicating that the CDR1as can play its role in sponging miR-7 (Hansen et al., 2013; Memczak et al., 2013). Plant circRNAs could target miRNAs to regulate gene expression at the post-transcriptional level (Tong et al., 2018; Xu et al., 2019; Hong et al., 2020; Ma et al., 2021). For example, in Arabidopsis, a circRNA named circ_At3g13990 has the potential to form a miR4239-5p-argonaut-circRNA ternary complex to downregulate the expression of target genes that have high expression in carpels and flowers (Frydrych Capelari et al., 2019). In *Brassica campestris*, it has been suggested that upregulation of a circRNA derived from the A02:23507399|23531438 locus could suppress a few miRNAs that target Bra002750 gene (involved in the regulation of several pollen development-related processes), resulting in up-regulation of this gene and production of abortive pollen (Liang et al., 2019). Also, a recent study using the CRISPR-Cas9 system in rice showed that a circRNA named Os06circ02797 could sponge and sequester Os-miR408 yielding faster-growing mutant seedlings with higher chlorophyll content compared with wild-type plants (Zhou et al., 2021).

**Figure 8.**
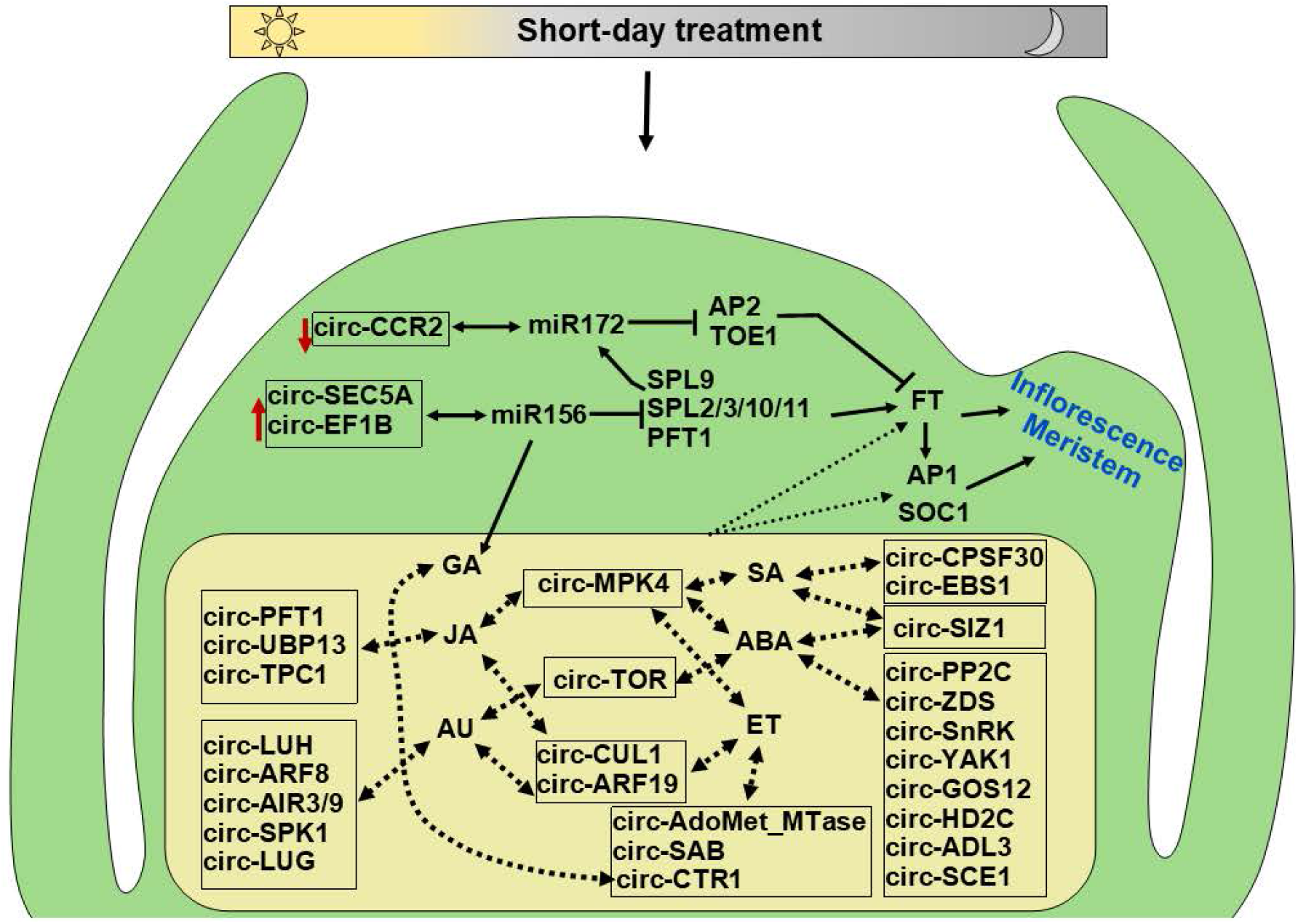
Proposed interaction of circRNAs with miR172 and miR156, and hormonal signaling pathway in shoot apical meristem of soybean. During the floral transition, the downregulation of circ-CCR2 increases the availability of miR172, leading to the downregulation of AP2 and TOE1 and the upregulation of FT. On the other hand, upregulation of circRNAs that sponge miR156, such as circ-SEC5A and circ-EF1B, decreases the availability of this miRNA, resulting in increased expression of SPLs and PFT1, which promotes expression of FT. Genes related to the hormonal signaling network showed circRNA production activity. These circRNAs might be involved regulating flowering genes and floral transition via interacting with hormonal signaling pathways through unknown mechanisms. Red arrows represent the upregulation or downregulation of specified circRNAs. Dashed arrows represent the possible interaction of circRNAs with hormonal signaling and flowering genes. Genomic position and properties of circRNAs are in Table S5.

Herein, we found that 38 differential-expressed circRNAs could potentially contain one to four putative binding sites for 39 miRNAs. Among the predicted miRNAs, we noted some important and well-studied miRNAs, such as miR156, miR159, miR169, miR171, miR172, miR319, and miR395 which all are involved in biological processes regulation during growth and development in plants including flowering (Vaucheret, 2009; Teotia and Tang, 2015; Dong et al., 2022). For example, the expression level of miR156 and miR172 regulates the developmental phase transition in plants (D’Ario et al., 2017; Song et al., 2019). At the early stages of plant growth, miR156 expresses at the highest level while miR172 has a low expression; as plants age, the expression level of miR156 decreases and the level of miR172 increases, resulting in the transition from vegetative to reproductive growth (Li et al., 2017). MiR156 targets and restricts genes related to stage transition and flowering time, such as SQUAMOSA PROMOTER BINDING PROTEIN-LIKE (SPL) genes, but miR172 restricts the expression level of APETALA 2 (AP2) genes which regulate floral transition, and flower meristem identity (Chen, 2004; D’Ario et al., 2017). Our analysis found four circRNAs that could target miR172 and four circRNAs with potential binding sites for miR156. One example in our analysis is circRNA Gm20:41203647-41204058 (circ-SEC5A), which is upregulated following the short-day photoperiod treatment. The up-regulation of this circRNA can potentially attenuate miR156 levels leading to up-regulation of the downstream flowering-related genes such as SPL9 to promote the floral transition in soybean SAM. SPL9 can activate the expression of miR172 and further promote flowering (Chen, 2004; Zhu et al., 2009). Based on our results and earlier findings, we suggest that circRNAs have the potential to play vital functions through the ceRNAs network during SAM development and floral transition in soybean.

### CircRNAs modulating hormonal pathways during floral transition

Since plant hormones regulate floral transition and flowering (Izawa, 2021), here we also investigated the expression of circRNAs from genes related to different hormonal pathways. The result revealed 40 genes with circRNA production activity with various hormonal pathways, including abscisic acid, auxin, jasmonic acid, salicylic acid, and ethylene. CircRNAs generated from these genes could potentially regulate biosynthesis or the signaling pathway of one or multiple hormones through a potentially complex interaction network leading to floral transition. Figure 8 illustrates a selected number of circRNAs that could interact with hormonal signaling pathways.

While gibberellin is one of the well-studied phytohormones in relation to flowering, the contribution and role of other hormones, such as abscisic acid, jasmonic acid, salicylic acid, and ethylene, have also been reported (Conti, 2017). For example, ABA is generally known as the drought-stress-induced hormone. Still, it’s positive regulatory effect on floral transition has been revealed, particularly via the activation of flowering genes such as CONSTANS (CO) and FT (Riboni et al., 2013; Riboni et al., 2016). Salicylic acid has also been reported to be involved in floral transition by regulating flowering genes such as FT, SOC1, and CO (Martínez et al., 2004). The transcription and regulation of multiple hormonal genes such as gibberellin, abscisic acid, jasmonic acid, and auxin have also been reported previously in soybean during floral transition in response to short-day treatment (Wong et al., 2009a; Wong et al., 2013a). Here in this study, we observed that genes related to hormonal pathways actively produce circRNAs in soybean SAM during the floral transition, especially genes related to abscisic acid and auxin. For instance, PROTEIN PHOSPHATASE TYPE 2C (PP2C) and SUCROSE-NON-FERMENTING-1-RELATED PROTEIN KINASE-2S (SnRK2) are the core components of ABA signaling in plants. We found that GmPP2C (Glyma.02g162300) and GmSnRK3.23 (Glyma.15g197300) produced circRNAs that might regulate the expression of ABA signaling components. We also found some circRNAs that could function in the biosynthesis processes of hormones. For example, ZETA-CAROTENE DESATURASE (ZDS) is an enzyme that works in the carotenoid biosynthesis pathway, a substrate for abscisic acid production in plants. Although the functional mechanisms of circRNAs derived from hormonal signaling pathway genes need further research, based on models proposed in plants, circRNAs could function in the regulation of their parent gene transcription via interacting with RNA polymerase II, form R-loop and alter the splicing of their locus by promoting exon skipping, affect translation by outcompeting mRNAs for binding the same proteins, or even directly bind mRNAs and increase their stability and translation (Conn et al., 2017; Liu and Chen, 2022).

### Functional Annotation analysis shows circRNAs can regulate meristem development and floral transition in short-day treated soybean SAM

Flowering in soybean is a complex process as each flowering gene found in Arabidopsis could have multiple orthologous copies in soybean (Jung et al., 2012; Wong et al., 2013b; Lin et al., 2021). Previous studies have shown that the gene expression in soybean SAM changes significantly during the floral transition with the potential function in several processes such as biosynthetic and metabolic process, regulation of cell size, and flower development (Wong et al., 2013a). Histone modifiers and RNAi-associated genes have also been reported to have high expression in soybean SAM during the floral transition (Liew et al., 2013).

To investigate the potential functions of circRNAs during floral transition in soybean SAM, we performed GO enrichment analysis on the parent genes of differential expressed circRNAs and their miRNA target genes. The GO terms belonged to biological process, molecular function, and cellular component categories (Table S10). For the parent genes of circRNAs, enriched GO terms were mainly related to protein modification and meristem development. For example, adaxial/abaxial axis specification, shoot apical meristem development, and specification of axis polarity were enriched GO terms related to meristem development (Figure 9A). To investigate the potential function of miRNA targets, we first predicted the target genes of 39 miRNAs that had binding sites on circRNAs, and the results showed that miRNAs could target more than 2300 unique genes (Table S11). The enriched GO terms of miRNA target genes belonged to various processes but were mainly related to reproduction and flowering. For instance, reproductive shoot system development, the developmental process involved in reproduction, and regulation of flower development were the GO terms that enriched the biological process category (Figure 9B; Table S12). These results suggest that circRNAs could be involved in diverse biological and molecular processes during floral transition in soybean SAM via direct regulatory functions or by targeting miRNAs in post-transcriptional regulation.

**Figure 9.**
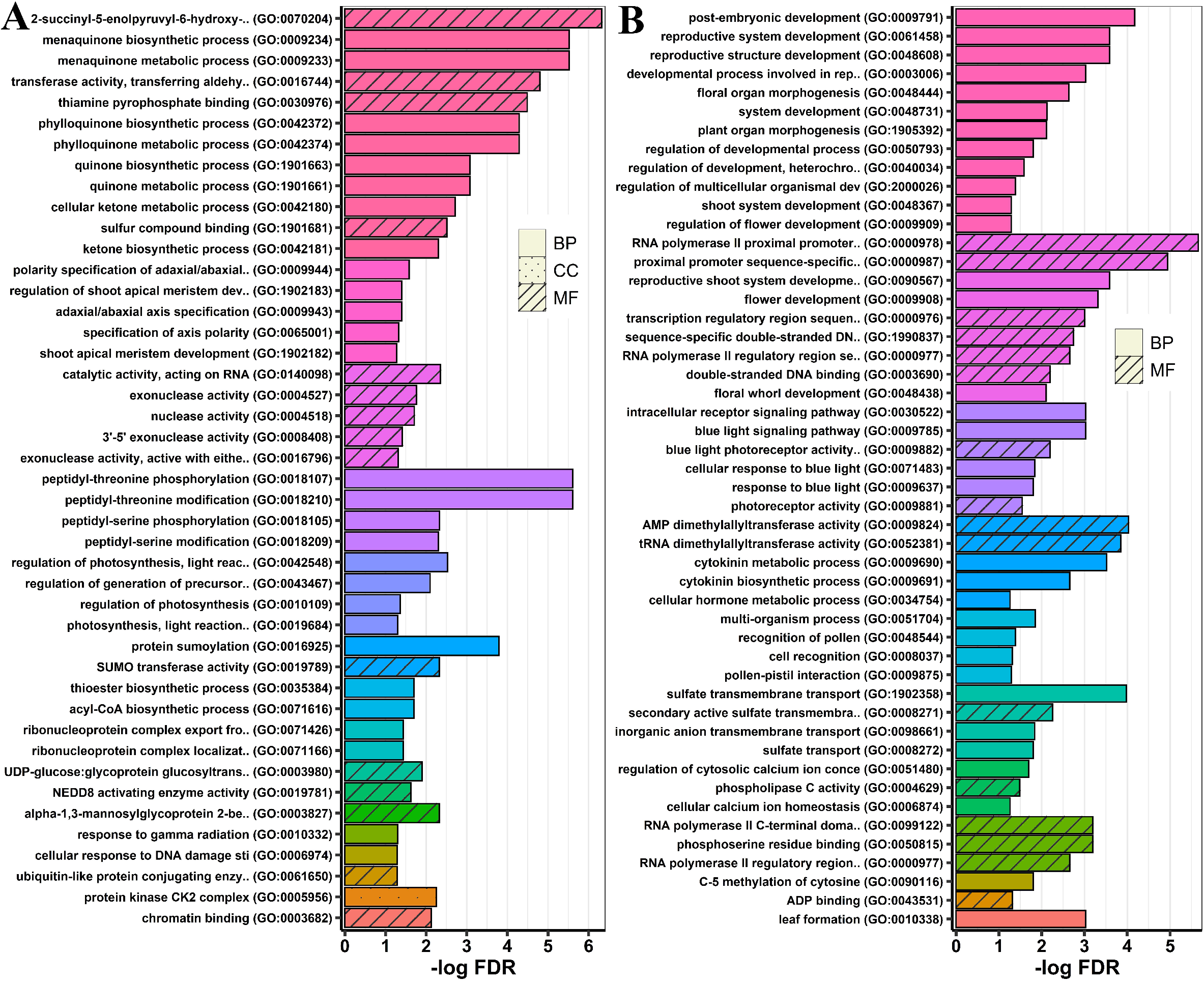
Functional enrichment analysis of circRNA parent genes (A) and their miRNA target (B). The enriched Gene Ontology (GO) terms for parent genes of circRNAs were mainly related to protein modification, photosynthesis, and meristem development, while for miRNA targets, GO terms were mainly related to flowering and reproduction. Bars with the same colour represent GO groups, as summarized by ClueGO.

In conclusion, our findings highlighted the expression profile of circRNAs in soybean SAM in response to short-day photoperiod treatment. We report 384 circRNAs, and most had differential expression during floral transition. circRNA-miRNA network analysis suggested the potential function of circRNAs as gene regulatory elements at the post-transcriptional level. Functional enrichment analysis revealed the host gene of circRNAs, and miRNA targets are mainly related to protein modification and meristem development, especially flowering. Our study also highlights the role of circRNAs in modulating the expression of hormonal signaling genes in regulating floral transition. Our results pave the way for further research to unlock the complexity of gene regulation during floral transition.

## Supplementary data

Supplementary Tables.xlsx: contains the annotation of all the identified circRNAs, and the results of related analysis. Supplementary Document.docx: contains gel electrophoreses photos and sequence information of validated circRNAs.

## Acknowledgments

This research was supported by Melbourne Bioinformatics at the University of Melbourne, project punim1093.

## Author Contributions

M.B.S. and P.L.B. designed and supervised the research; S.B. performed experiments, analyzed data, and prepared the figures, SB wrote the first draft, and all authors were involved in writing the manuscript.

## Data Availability

The RNA sequencing data is available in NCBI under accession number PRJNA893233.

